# Drastic genome reduction driven by parasitic lifestyle: Two complete genomes of endosymbiotic bacteria possibly hosted by a dinoflagellate

**DOI:** 10.1101/2025.01.03.631278

**Authors:** Takuro Nakayama, Ryo Harada, Akinori Yabuki, Mami Nomura, Kogiku Shiba, Kazuo Inaba, Yuji Inagaki

## Abstract

Bacteria with endosymbiotic lifestyles often exhibit dramatic genome reduction. While the reduction of genomes in intracellular symbionts of animals, including parasitic bacteria, has been extensively studied, less is known about such bacteria associated with single-celled eukaryotes. Here, we report the genomes of two novel gammaproteobacterial lineages, RS3 and XS4, identified as putative parasitic endosymbionts of the dinoflagellate *Citharistes regius*. Phylogenetic analyses suggest that RS3 and XS4 belong to the family Fastidiosibacteraceae within the order Beggiatoales, forming independent lineages therein. The genomes of RS3 and XS4 are 529 Kbp and 436 Kbp in size, respectively, revealing drastic genome reduction compared to related bacterial genomes. XS4, which has a particularly reduced genome with low GC content, uses an alternative genetic code, in which UGA assigned tryptophan. The small genomes of RS3 and XS4 encode a limited number of proteins, retaining only approximately 20% of the predicted ancestral proteome. Metabolic reconstruction suggests that RS3 and XS4 are energy parasites, heavily dependent on their host for essential metabolites. Furthermore, we found that the ancestor of both genomes likely acquired an ADP:ATP antiporter gene via horizontal gene transfer, an event that may have enabled their evolution as energy parasites by facilitating the acquisition of ATP from their host. These observations on novel bacteria with highly reduced genomes expand our understanding of the phylogenetic and genomic diversity of endosymbiotic bacteria in protists.

## Introduction

In natural environments, many bacteria establish symbiotic relationships with various eukaryotic organisms, including mutualistic, commensalistic, and parasitic interactions. The intracellular bacterial symbionts of arthropods are particularly well-documented: They provide metabolic complementation by supplying substances that the host cannot synthesize (e.g., specific amino acids) (McCutcheon and Moran, 2011), confer defensive capabilities against host predators (Oliver et al., 2003), or in some cases, manipulate host reproductive systems to facilitate their own transmission to the next generation (Katsuma et al., 2022). These symbiotic relationships have a significant impact on the evolution of these bacteria, particularly affecting their genomes. Bacteria that form intimate symbiotic relationships with eukaryotic hosts frequently undergo genome reduction, shedding genes that are unnecessary for their specialized lifestyle over evolutionary time. This can result in extremely small genomes, sometimes less than 1 Mbp in size (McCutcheon and Moran, 2011). Investigating these minimal genomes is crucial, not only to gain insight into specific mechanisms underlying each symbiosis but also to identify the essential gene sets required for bacterial survival (Koonin, 2003), which would provide valuable insights into fundamental biological processes.

To gain a comprehensive understanding of genome reduction in symbiotic bacteria, it is essential to investigate various symbiotic relationships hosted by a wide range of phylogenetic groups of eukaryotes. While previous studies have revealed that phylogenetically diverse bacteria engage in symbiotic relationships with eukaryotes, and have analyzed their genomes (McCutcheon and Moran, 2011; Moran et al., 2008), the majority of these studies have concentrated on symbiotic bacteria associated with animal hosts. However, advancements in molecular phylogenetics have demonstrated that multicellular organisms, including animals, represent only a small fraction of the overall eukaryotic diversity (Burki et al., 2020). The majority comprises single-celled eukaryotes, or protists, and these organisms are also frequently found to harbor symbiotic bacteria (Husnik et al., 2021; Nowack and Melkonian, 2010). Nevertheless, the phylogenetic and genomic diversity of the symbiotic bacteria in protists remains largely unexplored in comparison to those in animals (Husnik et al., 2021).

Dinoflagellates, belonging to Alveolata, represent a prominent group of protists thriving in various aquatic environments. These organisms exhibit substantial biodiversity, with some lineages known to have symbiotic bacteria (Foster and Zehr, 2019; Gavelis and Gile, 2018; Husnik et al., 2021), while genomic-level characterizations of these bacterial symbionts are scarce. Recently, genome analyses have been conducted on cyanobacteria symbiotic with marine Dinophysales dinoflagellates (Nakayama et al., 2024, 2019). Previous studies suggested the presence of additional, non-cyanobacterial symbionts within Dinophysales, including members of Gammaproteobacteria, Alphaproteobacteria, and Betaproteobacteria (Farnelid et al., 2010; Foster et al., 2006). However, only partial sequences of *nifH* and 16S rRNA genes have been obtained, which has left their metabolic characteristics largely unknown.

A recent study focused on and sequenced the genome of a cyanobacterial episymbiont of a Dinophysales dinoflagellate *Citharistes regius* (Nakayama et al., 2024). The cyanobacterial genome was recovered from a genome assembly generated from amplified DNA of a whole *C. regius* cell. Our re-analysis of this assembly also revealed additional genomic sequences representing putative endosymbiotic bacteria. Here we report on these genomic sequences and discuss the inferred evolutionary and metabolic characteristics of these bacteria. The analysis reveals that these bacteria, designated RS3 and XS4, have undergone extreme genome reduction, likely as an adaptation to their putative intracellular lifestyle and intimate association with the host dinoflagellate. They exhibit features of nutritional parasitism, relying heavily on their host for essential metabolites. Additionally, XS4 displays a unique genetic code alteration, further highlighting the evolutionary changes associated with its reduced genome and parasitic lifestyle.

## Materials and Methods

### Single-cell genome amplification, sequencing and assembly

Sampling of *Citharistes regius* and whole-genome amplification and genome assembly were performed as described in Nakayama et al. (2024). A preliminary annotation of the hybrid assembly of the amplified genomes was generated using the DFAST web service (Tanizawa et al., 2018, 2016). Predicted 16S rRNA gene sequences in the assembly were extracted and subsequently analyzed using the NCBI BLAST search service to determine their taxonomic origins to identify potential non-cyanobacterial genomes.

### Estimation of genetic codes and genome annotation

The preliminary genome annotation using the DFAST web service with default options revealed that, in the XS4 genome, several regions encoding proteins typically conserved in bacteria were interrupted by stop codons. This suggested that XS4 might utilize a genetic code different from that of typical bacteria. Therefore, we estimated the genetic codes of RS3 and XS4. This estimation focused on highly conserved amino acid positions within protein sequences conserved across bacterial lineages. We examined the codons used to encode these amino acids in the RS3 and XS4 genomes. The detailed procedure is as follows: First, we used the tBLASTn program to search the RS3 and XS4 genomes for regions likely to encode highly conserved, single-copy protein-coding genes. Sequence similarity searches of ribosomal RNA (rRNA) genes obtained from the preliminary annotation suggested that both RS3 and XS4 are closely related to *Fastidiosibacter lacustris* (genome accession number: GCF_003428155.1) among the bacterial genomes included in the Genome Taxonomy Database (GTDB) (Parks et al., 2022) r214. Therefore, we used the protein sequences of this genome as queries for tBLASTn. The target protein genes were 105 of the 120 genes encoding proteins used as phylogenetic markers in GTDB (bac120), detected from at least one of RS3 and XS4 (Supplementary Table S1). In parallel, we retrieved the aforementioned 105 proteins from 198 bacterial genome sequences representing diverse lineages included in GTDB r214 (Supplementary Table S1). Each set of 105 homologous protein sequences from the 198 bacteria was aligned using the L-INS-i method in MAFFT (version 7.490) (Katoh and Standley, 2013) to align evolutionarily homologous positions, resulting in 105 protein multiple alignments. From the amino acid positions in these alignments, we identified positions where 90% or more of the bacterial sequences had the same amino acid. We assumed that these highly conserved amino acids (hereafter HCAA) would also be utilized at the corresponding positions in the orthologous proteins of RS3 and XS4. Based on this assumption, we searched for sites corresponding to the HCAA positions within the homologous protein-coding sequences of the RS3 and XS4 genomes detected by tBLASTn, and investigated the codons used to encode these amino acids in each genome. The number of HCAA positions available for genetic code prediction in RS3 and XS4 were 10,707 and 10,214 sites, respectively. We counted the combinations of amino acid types at HCAA positions and the codons found in the RS3 and XS4 genomes, generating a 64 × 20 count matrix. This matrix was input into Weblogo3 (https://weblogo.threeplusone.com/create.cgi) (Crooks et al., 2004) to create sequence logos representing the amino acid frequencies at HCAA positions corresponding to each codon. The Weblogo3 options used were as follows: Units, probability; Composition, No adjustment for composition.

Genome annotation of RS3 and XS4 was performed using the DFAST web service (version 1.6.0) (Tanizawa et al., 2018, 2016). Analysis of the genetic code suggested that the proteins of the RS3 genome are encoded by the standard bacterial genetic code (NCBI’s translation table 11), while the proteins of the XS4 genome are encoded using a genetic code similar to that of *Mycoplasma* (NCBI’s translation table 4). Therefore, DFAST was run with genetic code option 11 for RS3 and genetic code option 4 for XS4.

### Phylogenotic analyses of 105 genes

We conducted two phylogenomic analyses: the first including the broad member of gammaproteobacterial species as operational taxonomic units (OTUs) and the second focusing specifically on species of the families Francisellaceae and Fastidiosibacteraceae as well as their close relatives. In both analyses, the multiple sequence alignments were generated based on the set of 120 phylogenetically informative marker genes (bac120) provided in GTDB r214. In the first analysis, we selected one representative genome from each order within Gammaproteobacteria and one representative genome from each genus within ‘Francisellales,’ the order employed within the GTDB. As outgroups, we included one representative genome each from Alphaproteobacteria, Zetaproteobacteria, and Magnetococcia. In cases where a lineage lacked a representative genome or had multiple representative genomes, the genome with the highest completeness was chosen. The bac120 homologs of RS3 and XS4 were manually selected from the results of BLASTp searches against the putative proteome of RS3 and XS4, using the bac120 dataset of the other genomes as a query. Among the bac120 dataset, 105 proteins were detected in at least one of RS3 and XS4 and were used for downstream analyses. The genomes and marker genes used in the analysis are listed in Supplementary Table S2. We aligned each of the 105 protein sequences using MAFFT (version 7.490) with the L-INS-i method and trimmed ambiguously aligned positions with BMGE (version 2.0) using default settings (Criscuolo and Gribaldo, 2010). The remaining sites of the 105 proteins were concatenated into a single alignment comprising 203 taxa and 33,898 amino acid positions. We inferred the maximum likelihood (ML) tree based on the alignment by using IQ-TREE (version 2.2.2.5) (Minh et al., 2020) under the LG+C10+F+I+G model, which was selected by ModelFinder of IQ-TREE (Kalyaanamoorthy et al., 2017) with ‘-m TEST -mset LG+C10’ option. The statistical support for each bipartition in the ML tree was calculated by 1000-replicate ultrafast bootstrap approximation (Hoang et al., 2018).

For the second phylogenomic analysis, we selected one genome from each species within Francisellaceae, Fastidiosibacteraceae, ‘Piscirickettsiales,’ and related environmental clades that were closely related to RS3 and XS4 in the first phylogenomic tree, totaling 44 genomes. The same 105 proteins used in the first analysis were retrieved from the 44 genomes (Supplementary Table S3), as well as the RS3 and XS4 genomes, and the sequences were aligned and trimmed in the same manner as in the first analysis. The 105 genes were concatenated into a single alignment comprising 46 taxa and 36,688 amino acid positions. The ML tree was inferred using IQ-TREE (version 2.2.6) under the LG+C60+F+I+G model, which was selected by ModelFinder with ‘-m TEST -mset LG+C60’ option. The statistical support for each bipartition in the ML tree was calculated by 100-replicate nonparametric bootstrap analysis with the ML tree used as the guide for estimating PMSF parameters (Wang et al., 2018).

### Phylogenetic analysis of ADP:ATP antiporter

We searched for the amino acid sequences of the ADP:ATP antiporter in the NCBI refseq database as of July 9, 2024, by BLASTp searches using the ADP:ATP antiporter sequences of RS3/XS4 as queries. We retrieved the subject sequences with E-values less than 1e-10, and divided the sequences into gammaproteobacterial, alphaproteobacterial, chlamydial, eukaryotic, and other sequences based on the lineages to which each sequence belongs. The redundancy within subject sequences of each phylogenetic group was removed by individual clustering analysis using CD-HIT (version 4.8.1) (Fu et al., 2012) with a 70% identity threshold. We aligned the representative sequences of all lineages and ADP:ATP antiporter sequences of RS3/XS4 using MAFFT (version 7.525) with the L-INS-i method. Ambiguously aligned positions were trimmed by trimAl (version 1.4) (Capella-Gutiérrez et al., 2009) with ‘-gt 0.85’ option. The final alignment comprised 229 sequences with 463 amino acid positions. We inferred the ML tree based on the alignment by using IQ-TREE (version 2.2.6) under the LG+C60+F+G model, which was selected by ModelFinder of IQ-TREE with ‘-m TEST -mset LG+C60’ option. The statistical support for each bipartition in the ML tree was calculated by 1000-replicate ultrafast bootstrap approximation.

### Orthologous protein analysis and functional annotation of proteins

Orthologous relationships between the proteins encoded in the RS3 and XS4 genomes and those of other Fastidiosibacteraceae bacteria were estimated using OrthoFinder (version 2.5.3) (Emms and Kelly, 2019). The OrthoFinder analysis included all proteins from RS3 and XS4, as well as those from eight species belonging to five genera of Fastidiosibacteraceae bacteria (Supplementary Table S4). To infer the metabolic functions associated with each orthologous protein group (orthogroup), one representative protein sequence was extracted from each orthogroup, creating a single protein set comprising a total of 2,935 sequences. This protein set was analyzed using KAAS (KEGG Automatic Annotation Server) (Moriya et al., 2007) to estimate the KO (KEGG Orthology) ID corresponding to each orthogroups (Kanehisa et al., 2016). The search program used in the KAAS analysis was BLAST, and the assignment method was BBH (bi-directional best hit). The metabolic function of each orthogroup was inferred based on the KEGG annotations associated with the assigned KO IDs.

## Results and Discussion

### Identification of circular genomes of RS3 and XS4

Nakayama et al. (2024) aimed to amplify the genome sequence of a cyanobacterium symbiotic with *Citharistes regius* by obtaining whole-genome amplification products from a whole *C. regius* cell (Nakayama et al., 2024). Given that the entire cell served as the template, the genomes of any organisms associated with this cell were theoretically amplified. Nakayama et al. (2024) focused on the symbiotic cyanobacteria and extracted only the cyanobacterial genome. In this study, we explored whether the amplification product contained any microbes other than cyanobacteria. As a result, we found two circular chromosomes showing homology to Gammaproteobacteria from an assembly of amplified genome using a single *C. regius* cell. The two gammaproteobacterial chromosomes have similar but clearly distinguishable ribosomal RNA gene sequences with an 89.95% identity, suggesting these chromosomes represent genomes of two phylogenetically related but separate bacteria (hereafter referred to as RS3 and XS4). The genome of RS3 was 529 Kbp in size with a GC content of 33%, whereas the genome of XS4 was 436 Kbp in size with a GC content of 28%.

In addition to these bacterial genomes, we also identified a potential complete archaeal genome sequence within the same assembly. This archaeal genome is exceptionally small and displays high sequence divergence compared to any known archaeal genomes. We consider that this genome originated from an archaeon representing a novel lineage. Therefore, discussing this archaeal genome alongside RS3 and XS4 within the same biological and evolutionary context is challenging and will be addressed in a separate publication.

### Genetic code in XS4

Preliminary annotation using the standard bacterial genetic code (NCBI’s translation table 11) revealed instances where open reading frames in the XS4 genome were interrupted by stop codons. This observation indicated the possibility of an alternative genetic code for XS4. Consequently, we estimated the genetic code employed in the putative protein-coding regions of RS3 and XS4 by comparing them with homologous proteins in other bacteria. We identified which codons on the genomes of RS3 and XS4 corresponded to conserved sites where identical amino acids are used in homologous proteins across various bacterial species (i.e., highly conserved amino acid site; Supplementary Figure S1). The results for the RS3 genome were consistent with translation using the standard bacterial genetic code. However, in XS4, the UGA codon (TGA in DNA), one of the three stop codons in the standard code, was located at sites where tryptophan (W) is conserved in other bacterial proteins (Supplementary Figure S1). This suggests that UGA codon is assigned for tryptophan in XS4. Such instances are also observed in organisms with low GC content in their genomes, including *Mycoplasma* and certain bacteria symbiotic with arthropod cells (Knight et al., 2001; McCutcheon and Moran, 2011). In the standard code, tryptophan is assigned by UGG, and the reassignment of UGA from translation termination to tryptophan is likely the result of selection pressures for low GC content in the genome (Knight et al., 2001; Osawa et al., 1992). We detected no sign of other deviations from the standard code in XS4. Consequently, the genetic code in the XS4 genome was inferred to be similar to that used in mycoplasmas, and the gene structure annotation of the XS4 genome was performed according to the *Mycoplasma* genetic code (NCBI’s translation table 4).

### Genomic features and phylogeny of RS3 and XS4

Genome annotation of the two genomes revealed 495 protein-coding genes in the RS3 genome and 426 in the XS4 genome. Each genome contained one copy of the 16S, 23S, and 5S rRNA genes, as well as 32 transfer RNA (tRNA) genes on RS3 and 31 tRNA genes on XS4, along with one transfer-messenger RNA (tmRNA) gene on both genomes (Figure 1). While the possibility of additional genomic elements such as plasmids cannot be excluded, the detection of rRNA genes and a complete set of tRNA genes corresponding to all 20 amino acids on both circular chromosomes strongly suggests that the genome sequences identified in this study represent the complete genomes of RS3 and XS4. Given that typical bacterial genomes are several million base pairs in size, the genomes of RS3 and XS4, which are 529 Kbp and 436 Kbp respectively, are remarkably compact.

**Figure 1.**
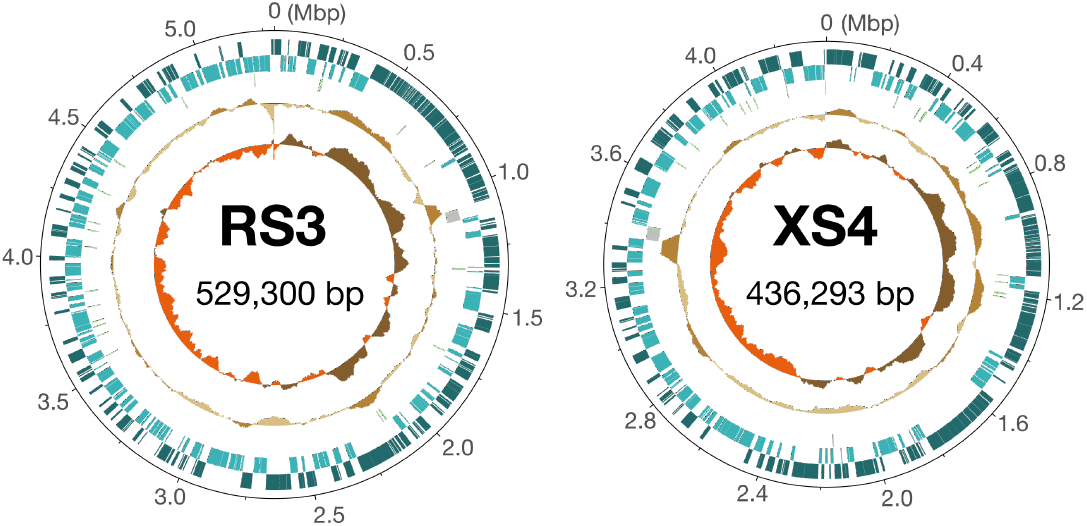
Genome maps of RS3 and XS4. For each map, the outer two circles indicate the positions of protein-coding genes on the plus and minus strands. The third circle shows positions of tRNA (green bars) and rRNA (grey bars) genes. The second innermost circle shows the relative G+C content and the innermost circle plots the GC skew.

Previous studies have detected *nifH* genes and 16S rRNA genes from bacteria other than cyanobacteria in Dinophysales dinoflagellates, including *Citharistes* sp. (Farnelid et al., 2010; Foster et al., 2006). The RS3 and XS4 genomes revealed in this study lack nitrogen fixation-related genes, including *nifH*. Thus, it is most likely that RS3 and XS4 have not been detected in the survey of the bacteria associated with Dinophysales cells using *nifH* sequences (Farnelid et al., 2010). Comparison with the 16S rRNA gene sequences obtained from Dinophysales cells in the previous study (Foster et al., 2006) was not possible, as the non-cyanobacterial sequences obtained were not deposited in public databases.

Subsequently, a multigene phylogenetic analysis was conducted to infer the phylogenetic positions of RS3 and XS4 among the bacterial diversity. From the preliminary sequence similarity search, it was suggested that both of these two bacteria are related to species in the families Fastidiosibacteraceae and Francisellaceae of Gammaproteobacteria. Firstly, we reconstructed a phylogenetic tree using genome sequences covering the diversity of Gammaproteobacteria, including Fastidiosi-bacteraceae and Francisellaceae, based on 105 protein sequences, which are highly conserved across bacterial lineages (Supplementary Figure S2). The topology of the tree suggested that both RS3 and XS4 are closely related to species within the Fastidiosibacteraceae. To gain further insight into the precise phylogenetic positions of RS3 and XS4, an additional phylogenetic analysis was conducted using a dataset including a more comprehensive sampling of fastidiosibacteracean and francisellacean species. Figure 2 shows the resulting ML phylogenetic tree, revealing that RS3 and XS4 constitute a monophyletic group within a cluster comprising Fastidiosibacteraceae bacteria. The lineage of RS3 and XS4 was shown to be a sister clade to the monophyletic group formed by the genera *Cysteiniphilum, Caedibacter*, and *Fastidiosibacter*. This monophyletic group, along with the clade comprising the genera *Fangia* and *Facilibium*, represents the monophyletic lineage of fastidiosibacteraceans. Additionally, a clade of Francisellaceae bacteria was inferred as a sister clade to the Fastidiosibacteraceae lineage. All the aforementioned phylogenetic relationships were robustly supported (bootstrap values of 100%).

**Figure 2.**
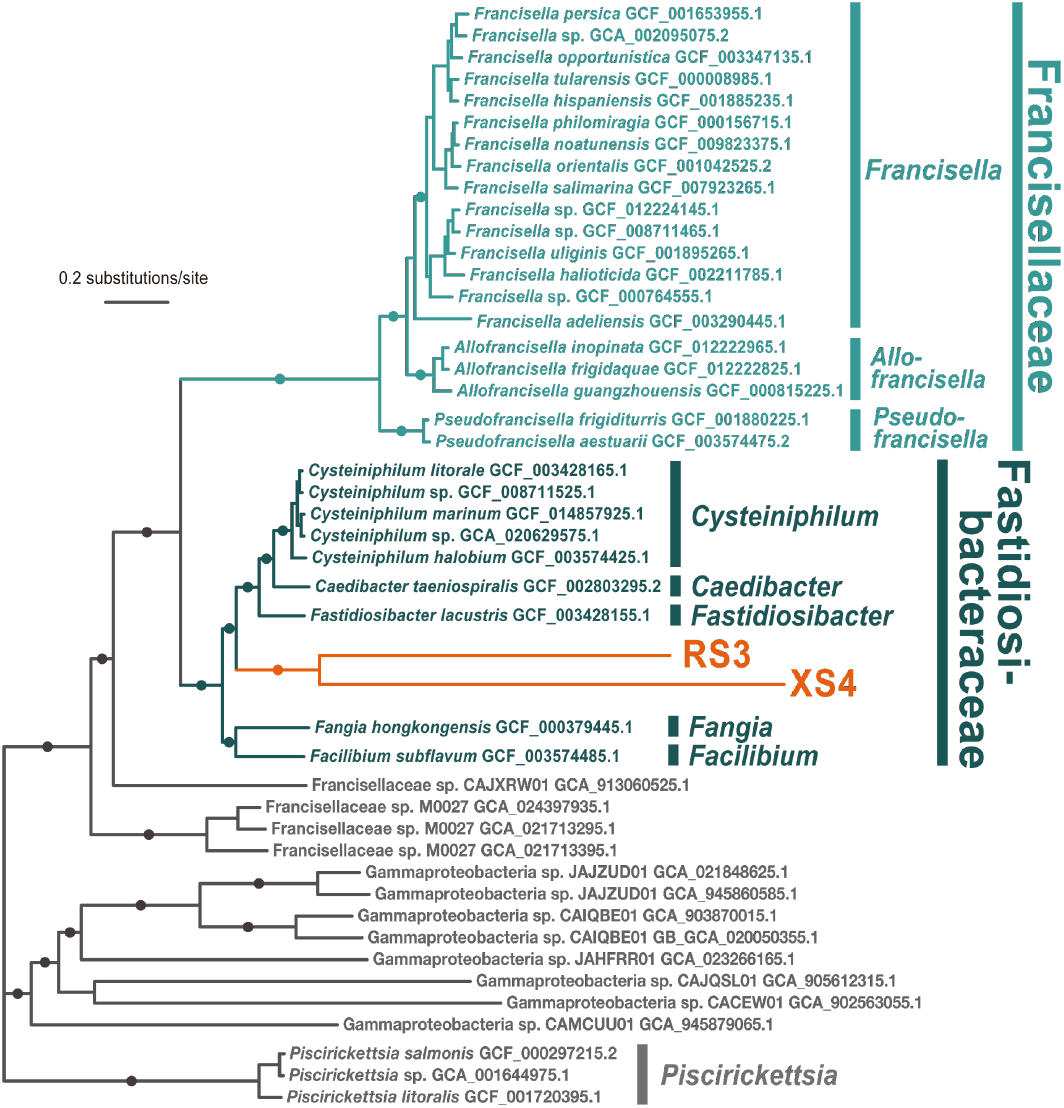
Maximum likelihood phylogenetic tree based on 105 protein sequences. The tree was inferred using IQ-Tree under the LG+C60+F+I+G model. Nonparametric bootstrap values (100 replicates) were mapped on the tree. The nodes marked with filled circles were supported with 100% bootstrap values. The alignment included 46 taxa and 36,688 amino acid positions.

The branches leading to RS3 and XS4 from the node connecting them are notably longer compared to those of other fastidiosibacteraceans, indicating a rapid evolutionary rate for these bacterial genomes. This topology of the tree is reminiscent of Long Branch Attraction (LBA) artifact, which is a methodological artifact where rapidly evolving lineages are incorrectly inferred to be closely related. However, given that the current analysis employed a mixture model accounting for site heterogeneity of amino acids frequencies, which is more robust to LBA (Quang et al., 2008), we consider this tree topology to represent the most plausible evolutionary hypothesis at present. The topology suggests that RS3 and XS4 are the closest relatives, diverging from a common ancestor shared with the genera *Cysteiniphilum, Caedibacter*, and *Fastidiosibacter*.

In the context of symbiotic relationships, including parasitism, *Caedibacter*, a member of the Fastidiosibacteraceae clade along with RS3 and XS4, is particularly noteworthy. The representative species *C. taeniospiralis* is an obligate cytoplasmic endosymbiont found in *Paramecium*, known for imparting toxic traits, referred to as ‘killer traits,’ to infected *Paramecium* against noninfected cells (Grosser et al., 2018; Kusch et al., 2000). Although most members of the Fastidiosibacteraceae have been discovered in the past two decades, research on *Paramecium* killer traits dates to the mid-20th century (Preer et al., 1974; Sonneborn, 1943), making *Caedibacter* the most well-known genus within this clade. The dinoflagellate *Citharistes regius*, the source of the genome amplification product from which RS3 and XS4 genomes were identified, is a protist belonging to Alveolata, as is *Paramecium*, implying a similar symbiotic association. Nevertheless, our phylogenetic analysis reveals that the *Caedibacter* and RS3/XS4 lineages are not closely related. In addition to *Caedibacter*, the marine fastidiosibacteracean genus *Cysteiniphilum*, which has been reported to parasitize humans (Xu et al., 2021), is also demonstrated not to be the closest relative of the RS3/XS4 lineage but to be a sister clade to *Caedibacter*.

### Genome reduction and codon reassignment

A comparative analysis of genome sizes based on the evolutionary relationships inferred by the phylogenomic tree reveals a notable reduction in genome size in RS3 and XS4. Figure 3 shows a comparison of genome size and GC content of each bacterial genome along with phylogenetic relationships: the genome sizes of *Fangia* and *Facilibium*, which represent the most basal branches of the Fastidiosibacteraceae lineage, are both slightly under 3 Mbp, and those of *Cysteiniphilum* and *Fastidiosibacter*, which are members of the sister clade of the RS3/XS4 lineage, range from 2-3 Mbp. In consideration of the genome sizes of the bacteria surrounding RS3 and XS4 in the phylogeny, it is likely that the ancestral bacteria of RS3 and XS4 possessed a genome of at least 2 Mbp. In contrast, the genome sizes of RS3 and XS4 are 529 Kbp and 436 Kbp respectively, indicating that their genomes have undergone severe reduction to less than a quarter of their ancestral size. Although the obligate cytoplasmic endosymbiont *Caedibacter taeniospiralis* is also known to possess a relatively compact genome, it is more than twice the size of the RS3/XS4 genomes.

**Figure 3.**
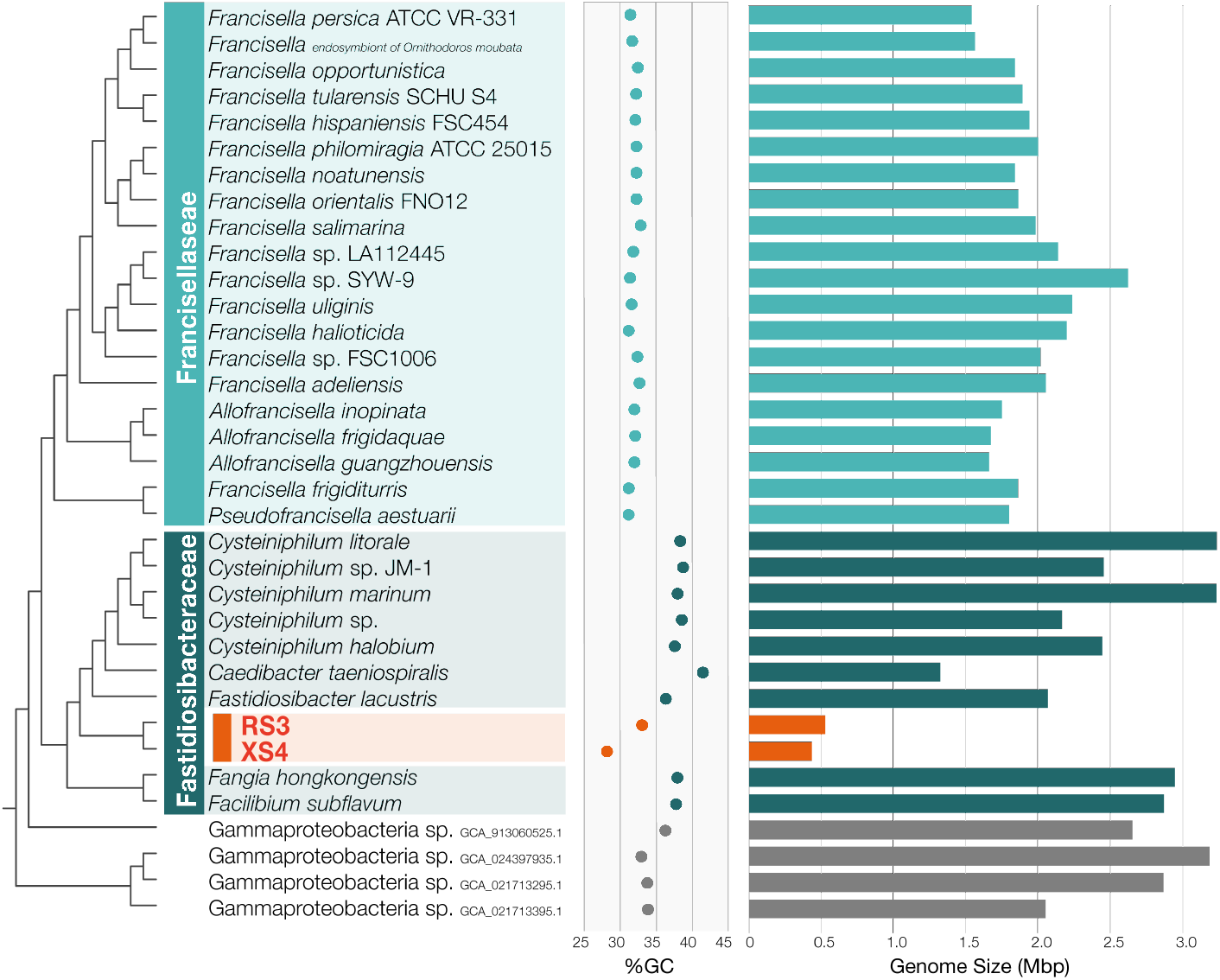
Comparison of size and GC content of the genomes of RS3, XS4, and their related bacteria. The cladogram on the left side shows the phylogenetic relationships as shown in Figure 2. The bar graph on the right displays the genome sizes, while the plot in the center indicates the GC content of each genome.

Another notable feature shared by RS3 and XS4 is their relatively low GC content (Figure 3). The GC content of the RS3 genome at 33.04% is low compared to other bacterial genomes of Fastidiosibacteraceae except for XS4. The XS4 genome exhibits an even lower GC content of 28.16%, which is less than those observed in genomes of Francisellaceae bacteria or those of fastidiosibacteracean genomes. Previous genomic studies have demonstrated a universal bias towards lower GC content in reduced genomes (McCutcheon and Moran, 2011), and a similar evolutionary trend appears to have occurred in the genomes of RS3 and XS4. The trend in GC content in reduced genomes is consistent with the observation that the XS4 genome, with its lower GC content, is more reduced in size than that of RS3.

In bacterial genomes that underwent severe size reduction and received a strong bias toward low GC content, UGA has often been reassigned from termination signal to tryptophan (Ohama et al., 2008). Interestingly, XS4 most likely uses the deviant genetic code, ‘UGA=W code,’ which assigns UGA to tryptophan, although RS3 uses the standard code (i.e., UGA as a termination signal). In bacteria with the standard code, two class-I release factors (i.e., RF1 and RF2), which promote the release of the nascent peptide from the ribosome responding to a stop codon at the A site, bear distinct codon specificities—RF1 recognized UAA and UAG, and RF2 recognized UAA and UGA (Caskey et al., 1969). Thus, there are two possibilities for RF2 in XS4. First, XS4 could have discarded RF2 during UGA reassignment, as RF1 is sufficient to cover UAA and UAG in this bacterium. Alternatively, RF2 persists but may have lost the codon specificity to UGA in XS4. The XS4 genome possesses the gene encoding RF2, and thus, this protein is anticipated to be UAA-specific. We compared the putative amino acid sequence of RF2 in XS4 to that of RF2 in RS3 (Note that the latter is supposed to recognize both UAA and UGA as termination signals). Unfortunately, the comparison provided no clue for the amino acid(s) that caused the change in codon specificity in RF2 in XS4 (Supplementary Figure S3). At least, the amino acids of RF2, which were shown to interact directly with UGA and UAA triplets in the crystal of the translation termination complex comprising the ribosome, a triplet (UAA or UGA), and RF2 (Korostelev et al., 2008; Weixlbaumer et al., 2008), are conserved in both XS4 and RS3 proteins.

Responding to the UGA reassignment, XS4 requires a tryptophan tRNA (tRNA^Trp^), of which anticodon can bind to UGA. We found only a single 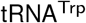 with the anticodon CCA 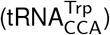 in both XS4 and RS3 genomes. While the CCA anticodon theoretically cannot translate UGA, as cytosine in the first anticodon position forms only a Watson-Crick pair with guanine in the third codon position, *Escherichia coli* 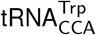 has been demonstrated experimentally that this tRNA can translate UGA codons with low efficiency (Hirsh, 1971; Weiner and Weber, 1971). Intriguingly, a recent work on a parasitic protist, *Blastocrithidia nonstop*, in which UGA is used as a tryptophan codon, demonstrated that the length of the anticodon stem of 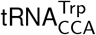 being critical for the translation of UGA (Kachale et al., 2023). In eukaryotes with the standard code (e.g., *Saccharomyces cerevisiae*), the anticodon stems of 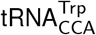 compose of five base pairs (5-bp AS) and binds solely to UGG. In contrast, *B. nonstop* has 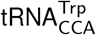 with the anticodon stem composed of four base pairs (4-bp AS) and the efficient translation of UGA by this non-canonical 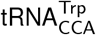 has been shown experimentally (Kachale et al., 2023). Surprisingly, the length of the anticodon stem in 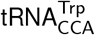 and the UGA assignments between XS4 and RS3 are consistent with this observation. We found 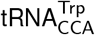 with a 4-bp AS in XS4 with the UGA=W code, while the same tRNA with a 5-bp AS was detected in RS3 with the standard code (Figure 4). Combined, we here propose that 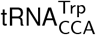 with a 4-bp AS recognizes UGA as a tryptophan codon, in addition to UGG, in XS4.

**Figure 4.**
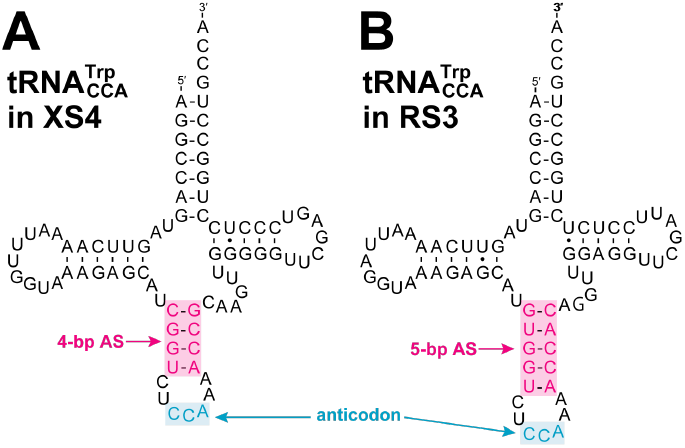
Predicted secondary structures of 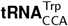. The anticodon stem of the tRNA in the XS4 genome (A) composed of four base pairs (4-bp AS) while that of the RS3 genome (B) have five base pairs (5-bp AS).

### Reductive evolution of metabolic capacities

Reflecting their small genome sizes, the genomes of RS3 and XS4 encode a correspondingly limited number of proteins: 495 and 426 protein-coding genes, respectively. In contrast, other Fastidiosibacteraceae bacteria, including species of *Cysteiniphilum, Facilibium, Fangia*, and *Fastidiosibacter*, have approximately 2,000-3,000 protein-coding genes in their genomes. Even the relatively compact genome of *Caedibacter taeniospiralis* contains 1,218 proteins. We conducted a comprehensive analysis of orthologous relationships between the proteins of RS3/XS4 and those of related bacteria to infer the evolution of the RS3 and XS4 proteomes in the light of the organismal phylogenetic relationship. Orthologous protein analysis revealed that nearly all proteins observed in the RS3/XS4 genomes (RS3: 480 out of 495 proteins, XS4: 408 out of 426 proteins) possess orthologous counterparts in their closely related bacterial species. By examining the proteomes of the phylogenetically related taxa, we inferred the minimal proteome of the common ancestor of RS3 and XS4. This ancestral proteome was defined as the set of proteins that satisfies two criteria below; The proteins found in: 1) either *Fangia* or *Facilibium*, representing the most basal branch of the Fastidiosibacteraceae clade; and 2) at least one member of the sister lineage to RS3/XS4, comprising *Cysteiniphilum, Fastidiosibacter*, or *Caedibacter*. The minimal ancestral proteome was estimated to comprise 2,151 proteins, with RS3 and XS4 retaining 477 and 402 of these proteins, respectively, which corresponds to only 22.1% and 18.6% of the ancestral proteome. Additionally, we assigned functional annotation using the KEGG Orthology (KO) database (Kanehisa et al., 2023) to each orthologous protein group in order to infer the metabolic capacity of RS3 and XS4 and the evolution of their proteomes. In comparison to the ancestral proteome, the majority of the metabolic functions lost were shared between RS3 and XS4 (Supplementary Table S5), suggesting these losses occurred at the stage of their common ancestor (Figure 5). Among the lost metabolic functions, transporters and two-component regulatory systems were the most affected functional categories, which are typically lost in symbiotic bacteria (McCutcheon and Moran, 2011; Shigenobu et al., 2000). Notably, both RS3 and XS4 lacked biosynthetic pathways for essential metabolites that are indispensable for independent growth. This is particularly evident in the biosynthesis of amino acids, where both bacteria have lost all major amino acid biosynthesis pathways. Additionally, both bacteria also lacked genes for the synthesis of pyrimidine bases, heme, the majority of vitamins, most of glycolytic enzymes, and half of TCA cycle enzyme genes. Moreover, XS4, with a more reductive genome than RS3, has lost additional metabolic pathways, including biosynthetic pathways for purine bases, chorismic acid, and riboflavin, and the entire set of the remaining TCA cycle enzyme genes, which are conserved in RS3. This extensive loss of metabolic functions indicates that RS3 and XS4 rely on their host for a significant proportion of metabolites required for their growth. Moreover, a distinguishing feature of RS3 and XS4 is the absence of biosynthetic pathways for peptidoglycan and lipopolysaccharide (LPS). While the morphology and symbiotic nature of these bacteria remain unclear, the loss of these key cell surface components suggests that both bacteria are likely intracellular symbionts (or parasites).

**Figure 5.**
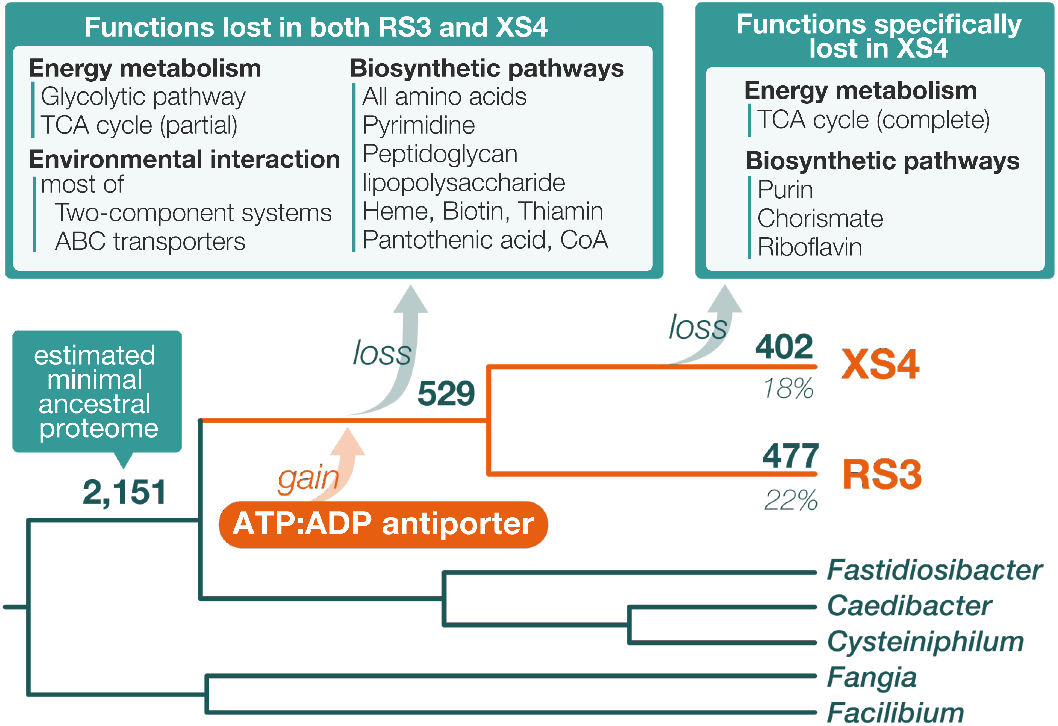
Predicted evolution of metabolic functions in the lineage of RS3 and XS4. Numbers above branches indicate the number of proteins derived from the minimal ancestral proteome (2,151 proteins), which was estimated based on the orthologous protein analysis. The percentages below the tips of the branches for RS3 and XS4 represent the proportion of proteins retained from the minimal ancestral proteome in each genome. Metabolic functions lost or gained during the evolutionary process of each bacterium are displayed.

A number of examples are known in which intracellular symbiotic bacteria synthesize and supply metabolic products that the host organism is unable to produce (McCutcheon and Moran, 2011). In such cases, both the host and the symbiont benefit from the relationship. Nevertheless, the relationship between RS3/XS4 and their host organisms is not necessarily analogous to the aforementioned cases. The lack of knowledge regarding the metabolic characteristics of the presumed host, *Citharistes regius*, makes it challenging to discuss the relationship in detail at present. However, RS3/XS4 have lost almost all pathways for synthesizing amino acids and vitamins, which makes it unlikely that they produce and supply the necessary substances for the host. On the other hand, it cannot be entirely ruled out that the residual biosynthetic pathways in RS3/XS4, such as those for folic acid and fatty acids, may still confer benefits to the host. This possibility can be examined once the metabolic functions of *C. regius* are elucidated in future studies. Regardless, it seems reasonable to assume that RS3/XS4 rely heavily on their host for the majority of the substances constituting their cells, characterizing these bacteria as primarily nutritional parasites. In this context, an intriguing protein-coding gene has been identified in both the genomes of these two bacteria. ADP:ATP antiporter (RS3CREG01_2930 and XS4CREG01_3620; KO ID: K03301), which was detected from the proteomes of RS3 and XS4, was not observed in any proteomes of Fastidiosibacteraceae bacteria examined in this study and was therefore not included in the estimated ancestral proteome of RS3/XS4. A phylogenetic analysis of the antiporter from various bacteria as well as those of RS3 and XS4 (Supplementary Figure S4) strongly supported the monophyletic relationship of the proteins from RS3 and XS4. Additionally, the RS3/XS4 clade was shown to be monophyletic with the alphaproteobacterial sequences, although the relationship was weakly supported. While homologous proteins were also identified from gammaproteobacterial species other than fastidiosibacteraceaens and francisellaceans, none of these were suggested to be closely related to the RS3/XS4 sequences (Supplementary Figure S4). The phylogenetic relationship indicates that the common ancestor of RS3/XS4 acquired the ADP:ATP antiporter gene through horizontal gene transfer, although the donor cannot yet be identified (Figure 5). It is noteworthy that several notorious intracellular parasitic bacteria, such as *Chlamydia* and *Rickettsia*, have been observed to engage in energy parasitism. These parasites have been shown to utilize the ADP:ATP antiporter as a key molecule for obtaining energy from their host organisms (Schmitz-Esser et al., 2004). Given these characteristics, it is hypothesized that the acquisition of this gene enabled RS3/XS4 to engage in energy parasitism.

## Conclusion

In this study, we report the genomes of RS3 and XS4, which were identified from amplified DNA obtained from the dinoflagellate *Citharistes regius* cell. RS3 and XS4 are regarded as novel symbiotic lineages belonging to the Fastidiosibacteraceae, Gammaproteobacteria. These bacteria possess highly reduced genomes (529 kbp for RS3 and 436 kbp for XS4), which are, to our knowledge, the smallest among the genomes sequenced at the chromosomal level within both Fastidiosibacteraceae and the encompassing order Beggiatoales (or Thiotrichales). Predicted proteomes of the two bacteria strongly suggest that RS3 and XS4 are adapted to an intracellular lifestyle. Furthermore, their markedly constrained biosynthetic capabilities indicate that these bacteria exist primarily as nutritional parasites within the host cells. The genomic characteristics of RS3 and XS4 revealed in this analysis not only expand our understanding of genome diversity within the Fastidiosibacteraceae but also contribute to the knowledge of the phylogenetic and genomic diversity of symbiotic bacteria in protists. Further exploration of bacteria symbiotic with protists is anticipated to yield even more insights.

## Acknowledgement

This work was supported by JSPS KAKENHI Grant Numbers JP16H06280, 20H03305, 18KK0203, 21K15131, 23K27226, 24K09587 22KJ0401 and BPI06050, as well as by the World Premier International Research Center Initiative (WPI Initiative), MEXT, Japan. R.H. was supported by JSPS Over-seas Research Fellowships. Computations in this study were partially performed on the NIG supercomputer at ROIS National Institute of Genetics. The manuscript file uploaded to bioR*χ*iv was generated using the LaTeX template adapted by Stephen Royle (https://github.com/quantixed) available at https://github.com/quantixed/manuscript-templates.

## Data availability

The annotated genome data of RS3 and XS4 have been deposited in DDBJ/GenBank/EMBL and will be made publicly available upon publication of the peer-reviewed article. Data related to the phylogenetic and orthologous protein analyses in this study are available on Zenodo (DOI: 10.5281/zenodo.14597766)

