## Supplementary Figure S4 for "Drastic genome reduction driven by parasitic lifestyle: Two complete genomes of endosymbiotic bacteria possibly hosted by a dinoflagellate"

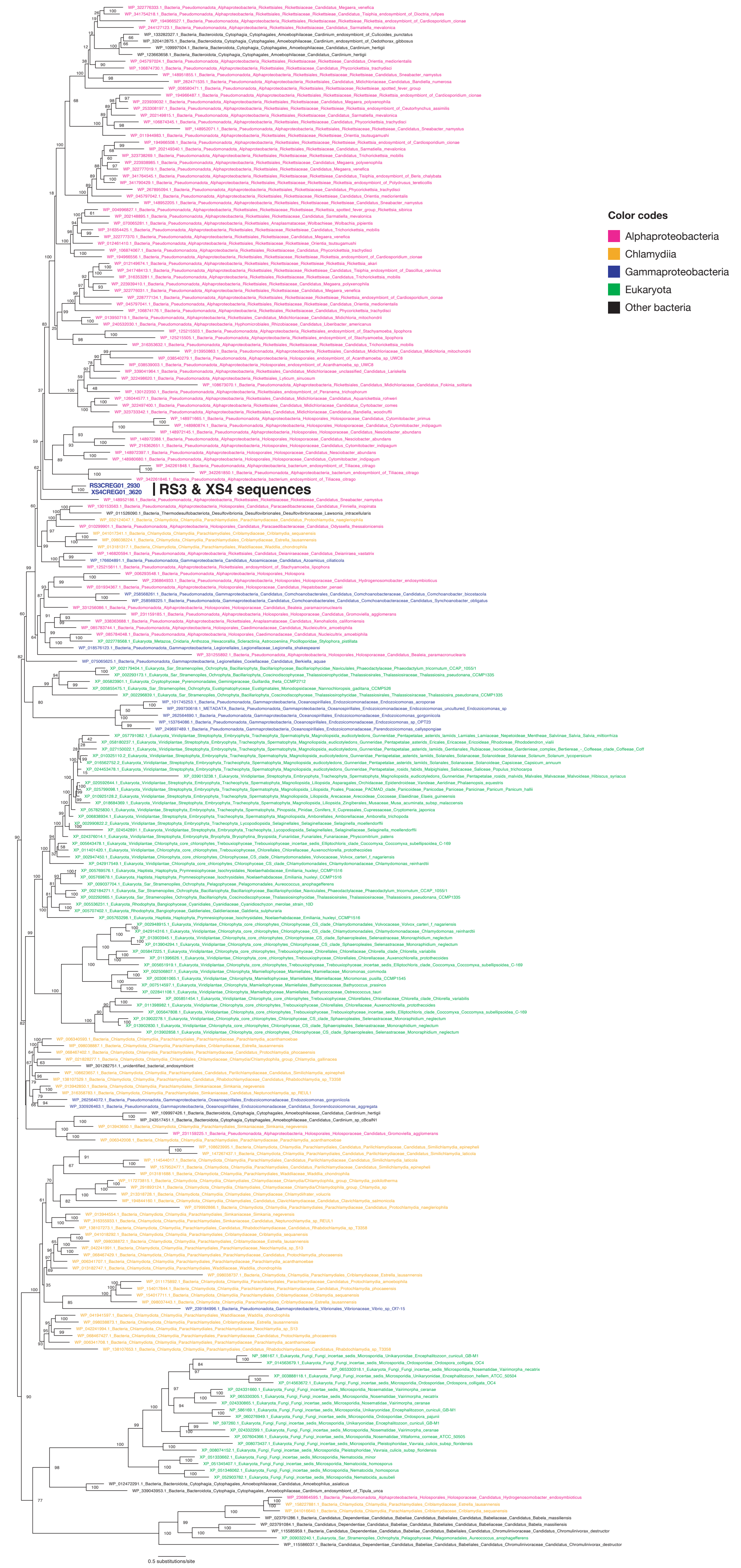


**Supplementary figure S4.** Maximum likelihood phylogenetic tree based on the ADP:ADP antiporter protein sequence alignment. The tree was inferred using IQ-Tree under the LG+C60+F+G model. The statistical support for each bipartition in the ML tree was calculated by 1000-replicate ultrafast bootstrap approximation. The alignment included 229 taxa and 463 amino acid positions.
