## Supplementary Figure S2 for "Drastic genome reduction driven by parasitic lifestyle: Two complete genomes of endosymbiotic bacteria possibly hosted by a dinoflagellate"

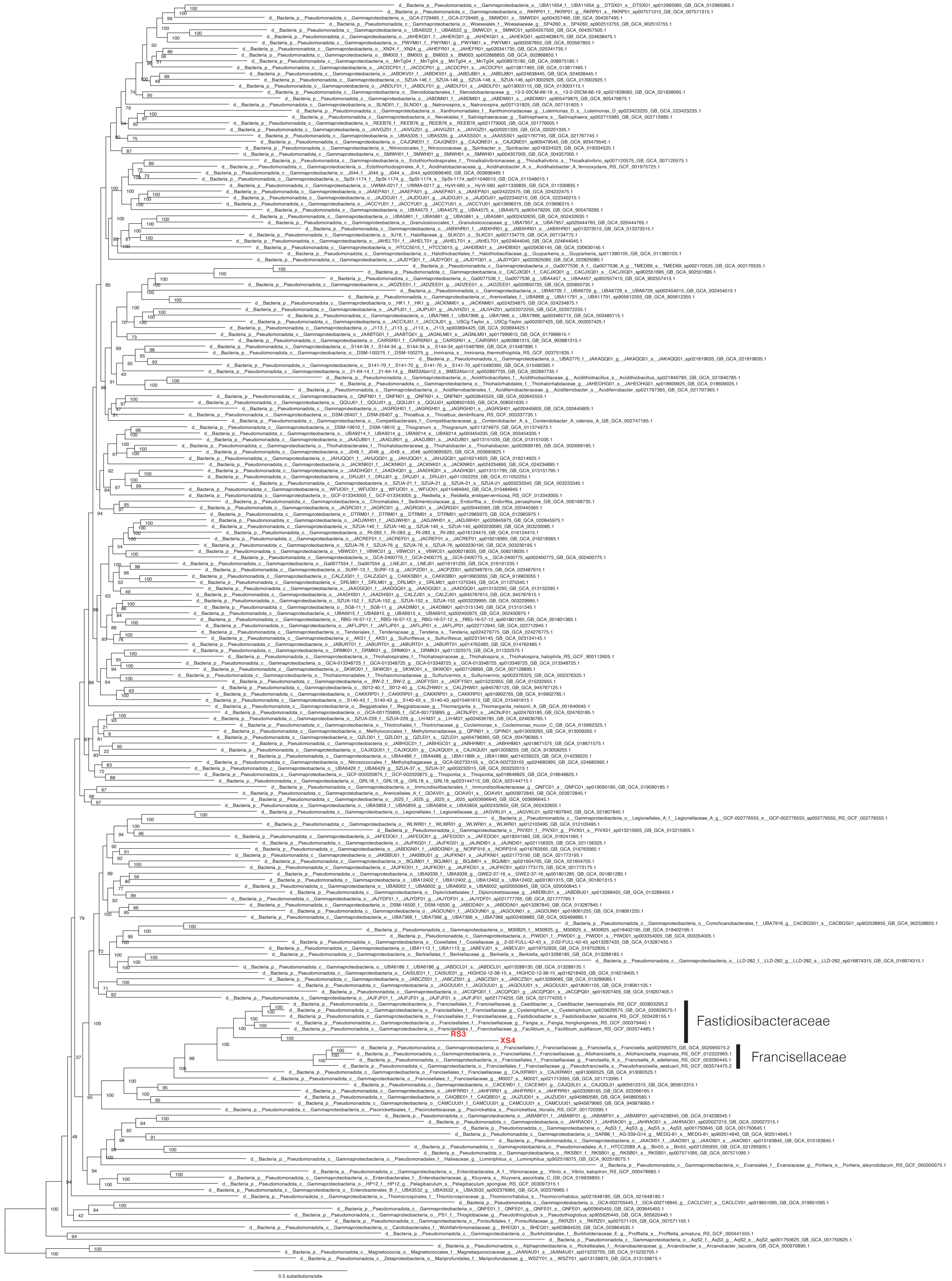


**Supplementary figure S2.** Maximum likelihood phylogenetic tree based on 105 protein sequences with broad taxon sampling. The tree was inferred using IQ-Tree under the LG+C10+F+I+G model. The statistical support for each bipartition in the ML tree was calculated by 1000-replicate ultrafast bootstrap approximation. The alignment included 203 taxa and 33,898 amino acid positions. The labeling of leaves is in accordance with the GTDB classification system.
