## Supplementary Figure S3 for "Drastic genome reduction driven by parasitic lifestyle: Two complete genomes of endosymbiotic bacteria possibly hosted by a dinoflagellate"

```
XS4 RF2      ------------------------------------------------------------
RS3 RF2      MKNEKNNKEFILQTKNNAYKSKLIDMKRRIIVIENISKIKTKEKRLEEIFKELESPKIWE
E.coli RF2   M------------FEINPVNNRIQDLTERSDVLRGYLDYDAKKERLEEVNAELEQPDVWN

XS4 RF2      ---------------------MNSLIDAVGTVSDLVSFSAQEKDDSIIEDILLEINNIKI
RS3 RF2      NKDYAKNINQEKINLQKTINNIKTLKNEVHEKLELLNFATENYEDFFFAEIIQEIHIFEK
E.coli RF2   EPERAQALGKERSSLEAVVDTLDQMKQGLEDVSGLLELAVEADDEETFNEAVAELDALEE

XS4 RF2      QLKRLELTCLFSNPLDSNNAFLDIQSGSGGTEAQDWAQILMRMYIRWGESHSFRVAITDI
RS3 RF2      KLKKFELVRKFSRNIDFNNAFLDIQSGSGGIEAQDWAQMLMRMYLKWGESHGFKTEITNI
E.coli RF2   KLAQLEFRRMFSGEYDSADCYLDIQAGSGGTEAQDWASMLERMYLRWAESRGFKTEIIEE

XS4 RF2      SHGEIAGIKSCTIHFMGKYSYGLLRTETGIHRLVRKSPFDSGNRRHTSFASVLSFPEIKD
RS3 RF2      KEGDVAGIKNCTISFKGEYAFCLLRTETGVHRLVRKSPFDASNRRHTSFASIFISPEINK
E.coli RF2   SEGEVAGIKSVTIKISGDYAYGWLRTETGVHRLVRKSPFDSGGRRHTSFSSAFVYPEVDD

XS4 RF2      DVSIEINFSDLRIDTYRASGAGGQHVNRTDSAVRVTHLPTNIVVQCQNNRSQHKNKDHAL
RS3 RF2      NIKVEINSSEIRVDTYKSSGAGGQHVNRTESAVRITHEPTSIVVQCQSDRSQHKNKEQAM
E.coli RF2   DIDIEINPADLRIDVYRASGAGGQHVNRTESAVRITHIPTGIVTQCQNDRSQHKNKDQAM

XS4 RF2      SQLKSKLYDLEVKKRNAKKQKLESEKLDITWGNQIRSYILDQSRIKDIRTNIEISNIQSV
RS3 RF2      KQLKSKLYEMKIREKENEKQKENINKSNISWGRHIRSYILDQSRVKDIRTGIENTNPQNV
E.coli RF2   KQMKAKLYELEMQKKNAEKQAMEDNKSDIGWGSQIRSYVLDDSRIKDLRTGVETRNTQAV

XS4 RF2      LDGKIDKFIYAALRL-L
RS3 RF2      LNGNLDEFIEESLKLNL
E.coli RF2   LDGSLDQFIEASLKAGL
```

**Supplementary figure S3.** Comparison among the RF2 amino acid sequences of *Escherichia coli*, XS4, and RS3. The amino acid residues highlighted in red were observed to interact with UAA/UGA triplet in the crystals of the translation termination complex. Two highly conserved motifs, which are responsible for codon recognition and promoting the hydrolysis of peptidyl-tRNA in the ribosome, are shaded.
