## Supplementary Figure S1 for "Drastic genome reduction driven by parasitic lifestyle: Two complete genomes of endosymbiotic bacteria possibly hosted by a dinoflagellate"

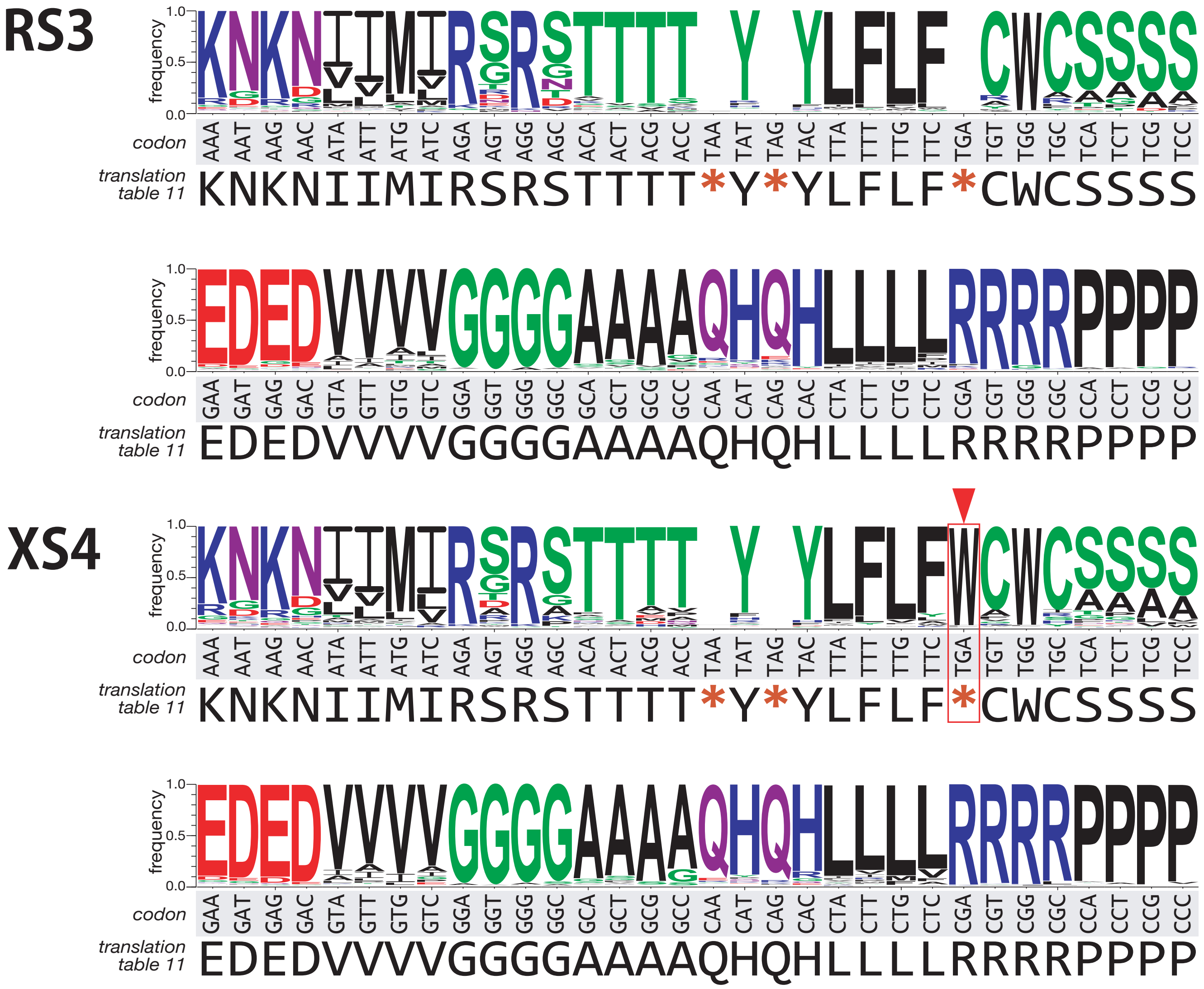


**Supplementary figure S1**. Relationship between codons in the RS3 and XS4 genomes and corresponding orthologous amino acids in other bacteria. The logoplot shows the proportion of amino acids observed at orthologous protein sites in bacterial sequences for each codon (DNA triplets) of putative protein-coding regions in the RS3 and XS4 genomes. The analysis focused on highly conserved sites where at least 90% of bacterial sequences, excluding RS3 and XS4, exhibited the same amino acid. the most frequently observed amino acid for each codon matched translations basedon NCBI's translation table 11. Similarly, in XS4, the results were largely consistent, although the amino acid site corresponding to TGA (UGA in mRNA) in other organisms predominantly exhibited tryptophan (W), suggesting that in XS4, TGA (UGA) codes for tryptophan instead of serving as a stop codon..
